# A search for snail-related answers to explain differences in response of *Schistosoma mansoni* to praziquantel treatment among responding and persistent hotspot villages along the Kenyan shore of Lake Victoria

**DOI:** 10.1101/394031

**Authors:** M. W. Mutuku, M. R. Laidemitt, B. R. Beechler, I. N. Mwangi, F. O. Otiato, E. L. Agola, H. Ochanda, B. Kamel, G. M. Mkoji, M. L. Steinauer, E. S. Loker

**Affiliations:** Centre for Biotechnology Research and Development, Kenya Medical Research Institute, Nairobi, Kenya; School of Biological Sciences, College of Biological and Physical Sciences, University of Nairobi, Nairobi, Kenya; Center for Evolutionary and Theoretical Immunology, Parasitology Division, Museum of Southwestern Biology, Department of Biology, University of New Mexico, Albuquerque, New Mexico, United States of America; College of Veterinary Medicine, Department of Biomedical Sciences, Oregon State University, Corvallis, Oregon, United States of America; Influenza Surveillance Program, Centers for Disease Control and Prevention, Nairobi, Kenya; Department of Basic Medical Sciences, Western University of Health Sciences, Lebanon, Oregon, United States of America

**Keywords:** *Schistosoma mansoni*, *Biomphalaria sudanica*, *Biomphalaria choanomphala*, Lake Victoria, persistent hotspots

## Abstract

Following a four-year annual praziquantel treatment campaign the resulting prevalence of *S. mansoni* was seen to differ among individual villages along the Kenyan shore of Lake Victoria. We have investigated possible inherent differences in snail-related aspects of transmission among such 10 villages, including six persistent hotspot (PHS) villages (≤30% reduction in prevalence following repeated treatments) located along the west-facing shore of the lake, and four PZQ-responding (RESP) villages (>30% prevalence reduction following repeated treatment) along Winam Gulf. When taking into account all sampling sites and times and water hyacinth presence/absence, shoreline-associated *B. sudanica* from PHS and RESP villages did not differ in relative abundance or prevalence of *S. mansoni* infection. Water hyacinth intrusions were associated with increased *B. sudanica* abundance. The deeper water snail *Biomphalaria choanomphala* was significantly more abundant in the PHS villages and prevalence of *S. mansoni* among villages both before and after control was positively correlated with *B. choanomphala* abundance. Worm recoveries from sentinel mice did not differ between PHS and RESP villages, and abundance of non-schistosome trematode species was not associated with *S. mansoni* abundance. *Biomphalaria choanomphala* provides an alternative, deepwater mode of transmission that may favor greater persistence of *S. mansoni* in PHS villages. As we found evidence for ongoing *S. mansoni* transmission in all 10 villages, we conclude conditions conducive for transmission and reinfection occur ubiquitously. This argues for an integrated, basin-wide plan for schistosomiasis control to counteract rapid reinfections facilitated by large snail populations and movements of infected people around the lake.

## INTRODUCTION

One of the most prevalent and persistent of the world’s neglected tropical diseases is human schistosomiasis.^1^ There is currently considerable momentum to bring schistosomiasis under control and to proceed to elimination efforts.^2–4^ An essential weapon in the elimination of schistosomiasis is the drug praziquantel (PZQ) which has been used extensively in a variety of control programs and has considerably reduced prevalence and intensity of infection.^5^ There is also a growing appreciation for the need for integrated control programs taking into account sanitation, provision of safe water, education, and acknowledging the fundamental role played by freshwater snails as vectors of the disease.^6,7^ The persistent success of schistosome parasites is owed substantially to their molluscan hosts which support the prolific production of human-infective cercariae, and that often exist in huge populations across a variety of freshwater habitats.

Africa harbors 90+% of the global burden of *S. mansoni*.^8^ In Kenya, there are three taxa of *Biomphalaria* snails that perpetuate transmission: 1) *B. pfeifferi*, whose distribution includes tributaries feeding Lake Victoria, and in small impoundments and both seasonal and perennial streams throughout the country, except in the tropical lowland belt along the coast; 2) *B. sudanica*, mainly found along the shores of Lake Victoria and Lake Jipe and their surrounding swamps; and 3) *B. choanomphala*, a deeper water inhabitant of Lake Victoria.^9,10^ The latter two taxa are likely members of the same species, and are frequently referred to as ecophenotypes or ecomorphs.^11–13^

Prevalence of intestinal schistosomiasis in Kenya is highest (>50%) in the Mwea irrigation scheme in central Kenya and in the Lake Victoria basin in western Kenya.^14^ Recent studies assessing the prevalence of *S. mansoni* infection in school children in the lake basin reported overall prevalences of 60.5% and 69%, with villages on lake islands having 2-fold higher prevalence rates than mainland villages.^15,16^ Schools closer to the lake also had higher prevalence rates than those further from the lake.^15^ Both higher prevalences and likelihood of reinfection following treatment would logically be associated with more frequent contact with lake water containing infected snails.^17^

Several chemotherapy-based initiatives have been undertaken to control schistosomiasis in the Lake Victoria region.^17–20^ Among them, the Schistosomiasis Consortium for Operational Research and Evaluation (SCORE) project implemented school-based and village-wide mass drug administration (MDA) in varying treatment regimens over 4 consecutive years (2011-2015) for 150 Kenyan villages situated on or near the eastern shore of Lake Victoria.^17^ Villages were randomized with respect to six different treatment strategies (study arms) which involved combinations of school-based or community-wide treatments, with 3 study arms receiving annual treatment and 3 receiving treatment twice over a four-year period. Regardless of study arm or age group of treated individuals, using spatial scan statistics a persistent hotspot area was identified in western Siaya County comprising west-facing villages along the open waters of the lake.^17^ Prevalence and intensity of infection did not decrease as much as it did in areas outside of the persistent hotspot area, the latter including villages along the more protected shores of Winam Gulf.^17^ Although this study could not clearly identify factors driving persistence of infection at hotspots, it reported an association between schistosome persistence and vegetation density (as measured by remote sensing) indicating a potential role for snail ecology.

Particular locations where the prevalence or intensity of schistosome infection does not fall to expected levels despite well-implemented, multiyear, MDA have been termed “persistent hotspots” or PHS, in contrast to responding villages (RESP) in which intensity and prevalence decline as expected. Why do some villages fail to respond to multi-annual treatment with PZQ? Several explanations can be offered including variability in efficacy of praziquantel among worm populations in different areas, levels of sanitation or availability of toilets, extent of drug coverage or compliance, availability of safe water supplies, extent of water contact, defecation behavior, or the extent to which they experience population movements such as resulting from fishing activities or proximity to ferry landings. ^19,22–25^

Conditions associated with snail vectors may also vary considerably including the nature of the snail species present, their abundance and local compatibility with schistosomes, longevity of infected snails and the rate at which they produce cercariae, and their patterns of distribution relative to water contact sites and anthropogenic disturbance.^22,26–33^ Shoreline and bottom conditions can also play important roles in efficacy of transmission.^9,22,24,32,34^

In addition to the shoreline-associated vegetation effects noted by Wiegand et al. as a possible factor associated with PHS, the dramatic presence and movements of large floating mats of the introduced water hyacinth, *Eichhornia crassipes*, are other factors in lake ecology that cannot be overlooked.^17^ Hyacinths have been suggested to provide a suitable environment for schistosome-susceptible snails and to facilitate their movements around the lake.^35^ Plummer noted that significantly more *B. sudanica* were recovered from experimental enclosures containing hyacinths than control enclosures lacking them, and Ofulla et al. also concluded that *B. sudanica* was significantly associated with hyacinths in Nyanza Gulf (now referred to as Winam Gulf), Kenya.^36,37^ Winam Gulf is one of the areas of the lake most affected by hyacinths, and four of the ten villages we investigated are found on the Gulf.^38^ Although hyacinth mats seem to favor snails in the short term, it has been noted that longer-term residence of hyacinth mats may be harmful to snails because of deficits in light penetration, water circulation, and oxygen availability, the latter due to decomposing hyacinths.^36^

Here we focus on the relationship of *Biomphalaria* snails to PZQ-based control efforts for *S. mansoni* in and around the Kenyan shoreline of Lake Victoria, one of the world’s major hyperendemic foci of human schistosomiasis. Upon the request of the SCORE program, during the two years immediately following the cessation of their MDA project, on four separate occasions, we surveyed *Biomphalaria* populations and their schistosome infection rates adjacent to 10 villages along the Kenyan shore of Lake Victoria with different degrees of responsiveness to treatment. In each of two collecting sites per village, we took note of the general condition of the habitat and searched for snails in shoreline-associated and deeper water habitats in the lake, as well as in smaller habitats adjacent to the lake. Building on the results of Wiegand et al., we investigated the hypotheses that PHS and RESP villages differ with respect to the species of vector snails present, relative snail abundance, likelihood that the snails are infected with schistosomes or other trematodes, and numbers of cercariae in the water as measured by infection rates of sentinel mice.^17^ All of these factors could contribute to persistently high prevalence in the face of control operations.

## MATERIALS AND METHODS

### Study sites

The study involved ten villages along the shores of Lake Victoria, western Kenya (Figure 1) that were part of the Wiegand et al., study already mentioned.^17^ The villages had annual school-based mass drug administration (MDA) for 4 consecutive years (Table 1). Following one of the definitions of Kittur et al., six of the villages (Minya, Agok, Migiro, Miyandhe, Kanyibok, Usenge) were persistent hotspots (PHS), locations where the absolute change in *S. mansoni* prevalence from the beginning of the control program to the end was ≤30%.^21^ By contrast, for the four remaining villages (Kotieno, Seka Dok, St. Douglas Weta, Mumbo) recorded a drop in prevalence >30% and are considered responding (RESP) villages. In each village, two shoreline habitats were identified where there was evidence of human-water contact activities, and these were established as our sampling sites. Each of the 20 sampling sites was visited four times (April 2016, August 2016, January 2017 and May 2017).

**Figure 1:**
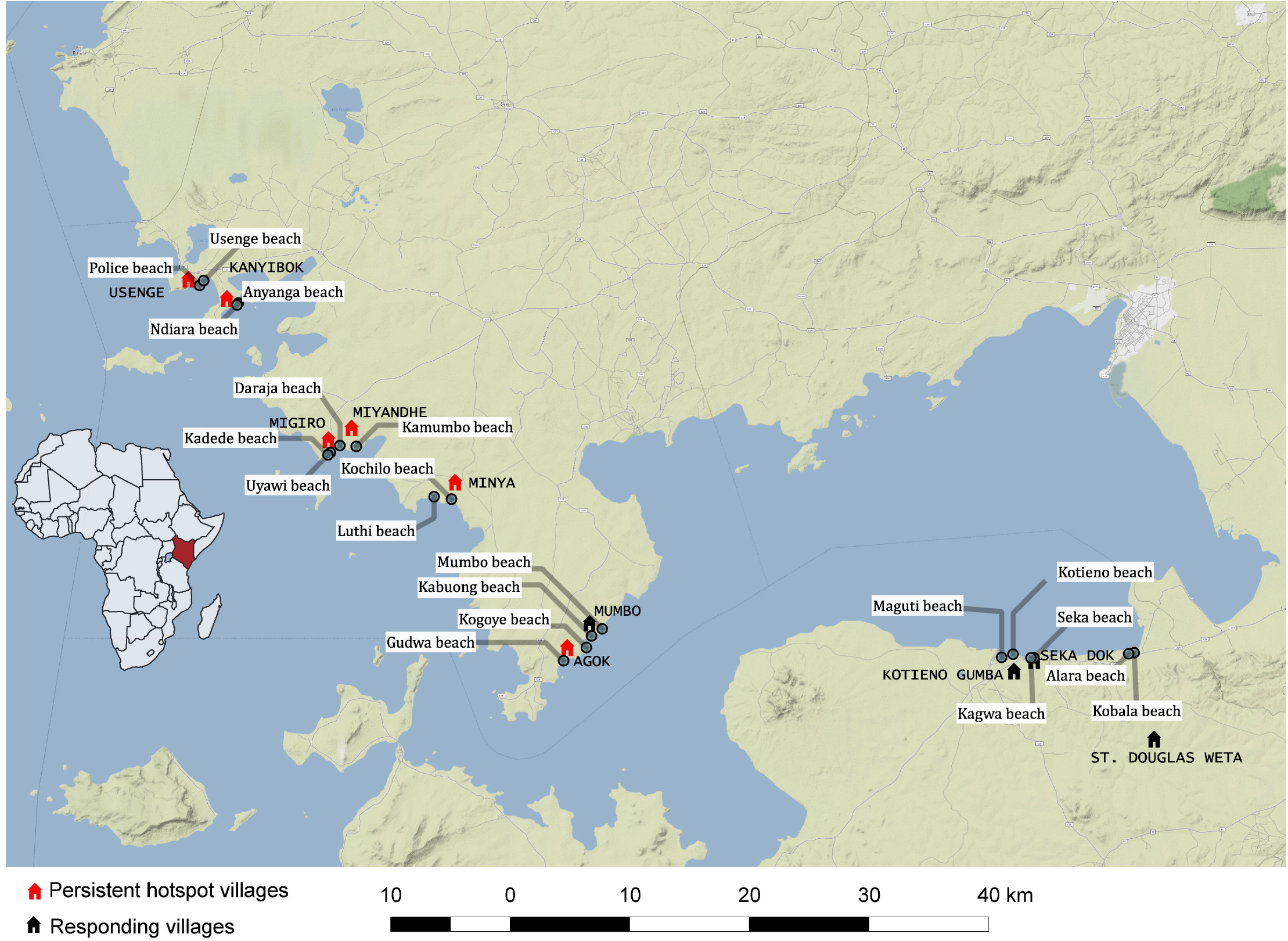
Map showing position of study villages along the shore of Lake Victoria, Kenya. The map was constructed using QGIS version 2.18.22 and 3.2.1, the basemap layer was added from the standard terrain tiles provided from http://maps.stamen.com through the OpenLayers function. The tiles are © Stamen Design, under a creative commons attribution (CC BY 3.0) license. The data for the locations were obtained manually using cellular phone GPS functionality according to the WGS1984 mercator projection.

**Table 1:** *Schistosoma mansoni* prevalence in schools within the study villages showing response to praziquantel treatment following four years of Mass Drug Administration. Data acquired from and shown with permission of the SCORE project. Sample sizes shown in parentheses.

| SCHOOL | Prevalence (%)<br>Year 1 (2011) | Prevalence<br>(%) Year 2<br>(2012) | Prevalence<br>(%) Year 3<br>(2013) | Prevalence<br>(%) Year 4<br>(2014) | Prevalence<br>(%) Year 5<br>(2015) | Absolute<br>Prevalence<br>Change<br>Year 1 Minus |
| --- | --- | --- | --- | --- | --- | --- |

|  |  |  |  |  |  | Year 5 as % |
| --- | --- | --- | --- | --- | --- | --- |
|  | **Persistent Hotspot Villages** |  |  |  |  |  |
| MINYA | 68.6 (n=97) | 77(n=100) | 75(n-97) | 74(n-92) | 82(n-97) | +13.4 |
| AGOK | 60.5(n-99) | 59.8(n-93) | 60.2(n-97) | 67(n-86) | 39(n-91) | -21.5 |
| MIGIRO | 69.4(100) | 76.7(n-94) | 49.2(n-87) | 72(n-87) | 48(n-88) | -21.4 |
| MIYANDHE | 78.6(86) | 70.5(n-91) | 71.4(n-87) | 73.5(n-92) | 54(n-94) | -24.6 |
| KANYIBOK | 91.8(100) | 93(n-96) | 83.7(n-98) | 84(n-94) | 83(n-100) | -8.8 |
| USENGE | 69.4(100) | 68(n-100) | 65(n-100) | 46.5(n-100) | 65(n-100) | -4.4 |
|  | **Responding Villages** |  |  |  |  |  |
| KOTIENO | 52.9(87) | 40.9(n-92) | 19(n-93) | 16(n-84) | 12(n-88) | -40.9 |
| SEKA DOK | 81.6(100) | 73.2(n-97) | 55.8(n-91) | 48(n-90) | 19(n-96) | -62.6 |
| ST.DOUGLA<br>S WETA | 55.2(100) | 27.4(n-94) | 24(n-93) | 19(n-87) | 19(n-91) | -36.2 |
| MUMBO | 57.7(77) | 30(n-74) | 11.5(n-78) | 15.6(n-79) | 5.5(n-77) | -52.5 |

### Snail Sampling

Two methods were used to collect snails from each of the 20 designated sampling sites: scooping from the shore and dredging from a boat. From the shore, two experienced collectors from our team scooped snails for 30 minutes per sampling site using long-handled scoops (steel sieve with a mesh size of 2 × 2 mm, supported on an iron frame). Shoreline habitats were typical for Lake Victoria consisting of emergent and submerged vegetation that was swept by scoops for snails. Snails were also collected offshore by passing a dredge (0.75 m long and 0.4 m wide with attached sieve, 2 × 2 mm mesh size) along the bottom. At each sampling site dredge hauls covering a combined length of 150 meters was made, beginning at one meter depth and extending perpendicular to the shore to 10m depth. Live snails were picked from the dredged material. All snails were taken to the laboratory at KEMRI, Center for Global Health Research, Kisumu where they were sorted into species based on shell morphology characteristics, using standard taxonomic identification keys, counted and screened.^39^

### Snail screening

Each snail was placed in an individual well of a 24-well plastic culture plate containing 1 ml of aged de-chlorinated tap water. The plate was placed in indirect sunlight for 2 hours between 10:00-13:00hr. Individual wells were then examined with the aid of a dissecting microscope for presence of cercariae. Using standard identification keys, cercariae were identified to basic taxonomic groups.^40^ Non-shedding snails were maintained in the laboratory for four weeks and screened again to enable detection of snails that harbored pre-patent infections at the time of initial collection.^31^ Some cercariae were further identified with molecular methods as parts of another study.^41^ Additionally, samples of mammalian schistosome cercariae recovered from *B. sudanica* (each sample consisting of 4 cercariae) were extracted (QIAamp DNA Micro Kit, to elution volume of 45μl) and primers for the mitochondrial NADH dehydrogenase subunit 5 gene (*nad5*) used for amplification, following which bands were visualized on a gel (302 bp expected band size for *S. mansoni* and 800 bp for the related schistosome *S. rodhaini*) as a check for the identity of the schistosome recovered.^28^

### Sentinel mice exposures

Mice were exposed to lake water at each near shore sampling site in floating cages that ensured the mice were exposed to a depth of no more than 5 mm. Each cage was made from a plastic container measuring 20 × 14 × 7 cm with the bottom of the container replaced with wire mesh with openings measuring 5 × 5 mm. Styrofoam blocks were attached to the plastic container to insure the cage floated at the desired position. Lids of the containers were dark to provide shading and were perforated with 5 mm diameter holes to allow air circulation. The abdomen of each mouse was shaved to facilitate cercariae penetration. For each of the shoreline sampling sites, five cages each with 2 mice were tethered adjacent to locations where snails were collected (total of 20 mice per village per sampling time). Mice were in contact with the water for 3 hours (11:00 hr – 14:00 hr), an interval when shedding of cercariae by *S. mansoni*-infected snails is expected to be at its peak.^42^ Previous experience under lakeside conditions indicated that a three-hour period of exposure did not jeopardize survival of the mice, which was confirmed by the high survival rates of the mice retrieved. Mice from each sampling site were then marked with an identifying ear tag and maintained together until perfused at eight weeks post-exposure. Recovered worms were counted and tabulated with respect to village, sampling site and sampling time.^43^

### Ethics statement

All experiments involving mice were approved by institutional animal care and use committees (IACUCs) at the Kenya Medical Research Institute (Protocol KEMRI/ACUC/03.10.15) and at the University of New Mexico (Animal Welfare Assurance # D16-00565 (A4023-01, expiration 8/29/2021). All protocols and practices for the handling and manipulation of mice were in accordance with the guidelines of the American Veterinary Medical Association (AVMA) for humane treatment of laboratory animals.

### Statistical analysis

To determine if PHS sites differed from RESP sites in snail-related factors, we compared them with respect to relative abundance of each snail (*B. sudanica* or *B. choanomphala*), the prevalence of infected snails, and the number of adult schistosome worms collected from the sentinel mice (as a measure of the force of infection). Collection sites were nested into village and were compared across the four time points. Because of the potential effects of large rafts of floating water hyacinths that temporarily occupy some sites, we included the presence or absence of hyacinths in the models.

#### Snail relative abundance

To determine whether relative abundance of either *B. sudanica* or *B. choanomphala* differed between PHS and RESP villages, between sites with hyacinths and no hyacinths, or over time, we performed generalized linear mixed models in R using the package glmmTMB, with the random effect of village and using the family negative binomial 1 (nbinom1).^44^ Prior to settling upon this model structure we also explored a model that included a nested random effect of sample site within village but the model fit was not significantly better (anova, p>0.05) so we report the reduced model of only village as a random effect.

Formula: glmmTMB(Abundance of either snail ~ Responder/Not + Hyacinth Y/N + timepoint + (1 | School),family=nbinom1(link=log))

#### Prevalence of infected snails

To determine whether the likelihood of detecting an infected snail was different in a RESP vs. PHS village, or in sites with and without hyacinths we performed a generalized linear mixed model using R with the packages lmerTest and lme4 with the random effect of village and the family of binomial (logit link).^45^ We also included the offset term of number of snails found, rescaled to be on the same scale as the binomial variables by dividing by 2 standard deviations.^46^ This is because there is a strong positive correlation between the likelihood of finding an infected snail and the total number of snails found (est=0.592+/-0.148, p=0.00005, statistics reported from the same model as described above but with total number of snails found included as a main effect rather than an offset). We analyzed only data for *B. sudanica* because the number of infected *B. choanomphala* was prohibitively small.

Formula: glmer(smansoni.presence Y/N~ Responder/Not+ Hyacinth Y/N+ (1| timepoint)+(offset(rescaledtotalsudanicacount))+(1| School),family=binomial(link=“logit”))

#### Sentinel mice

To evaluate whether infection prevalence in the sentinel mice differed between PHS and RESP villages, between sites with and without hyacinths and between time points we performed a generalized linear mixed model in R using the package glmmTMB with a random effect of village, and family of negative binomial 1. We also included the term of total snails found at each site to account for snail relative abundance, however this variable was rescaled by dividing by 2 standard deviations to place it on the same scale as the binomial variables.

Formula: glmmTMB(total.number.of.schistosomes.in.mice ~ Responder/Not + Time + total.biomphalaria.snails + (1 | Village), family=nbinom1(link=”log))

We also evaluated whether the likelihood of finding a schistosome positive sentinel mouse differed between PHS and RESP villages, between sites with and without hyacinths and between time points. The only difference between this model and the one described above is that the dependent variable is binomial (infected mouse found or not), rather than the burden within the infected mouse. For this we used a model structure similar to the prevalence of infected snails. We performed a generalized linear mixed model with family of binomial (link=logit) and the random effect of village. Once again we accounted for the total number of snails collected, but rescaled it to be on the same scale as the binomial variables.^46^

Formula: glmer(smansoni.presenceinmice Y/N~ Responder/Not+ Hyacinth Y/N+ (1| timepoint)+(offset(rescaledtotalbiomphalariacount))+(1| Village),family=binomial(link=“logit”))

#### Correlation of human prevalence and snail density

To determine if the abundance of snails at a particular site was associated with schistosome infection rates in humans, we correlated the total number of snails collected at a village (summed between collecting sites and all 4 time points) with both the initial and the final prevalence in humans at the corresponding village using Pearson’s correlation in Graphpad Prism 7.0. We performed separate analyses for each snail species.

#### Biodiversity

To assess whether the likelihood of detecting a schistosome-infected *B. sudanica* at a site correlated with the presence of other digenetic trematodes in the snails, we included each of four commonly collected cercarial types (echinostome, strigeid, xiphidiocercariae, and amphistome) as a main effect in the same model as described above in the prevalence of infected snails.

Formula: glmer(smansoni.presence Y/N~ Responder/Not+ Hyacinth Y/N+amphistomes Y/N+xiphido Y/N, echinostomes Y/N, strigieds Y/N + (1| timepoint) + (offset(rescaledtotalsudanicacount)) + (1| village),family=binomial(link=“logit”))

## RESULTS

### Descriptive overview

A total of 12,156 snails (10,249 *B. sudanica* and 1,907 *B. choanomphala*) were collected from the study sites during four collection periods (Table 2, Figure 2). No *B. pfeifferi* were found in the lake habitats sampled. We also searched small streams or impoundments near the villages and found them to be muddy, prone to drying and without *Biomphalaria* snails. At some times and collecting sites, hyacinth mats covered the water surface and were so dense as to preclude dredging (Figure 2). *Biomphalaria sudanica* was recovered along the shoreline from all 20 sampling locations and *B. choanomphala* was dredged from deeper water at all 12 sampling locations in PHS villages whereas Mumbo was the only RESP village from which *B. choanomphala* was found (Table 2). Of note is that Mumbo is located closer to the PHS villages than the remaining three RESP villages (Figure 1). *Biomphalaria choanomphala* was also sometimes found while scooping from shoreline collecting sites in three PHS locations, Kanyibok, Migoro and Usenge. At Usenge Beach, the shoreline *B. choanomphala* population persisted for at least five months on submerged vegetation along with another snail customarily recovered from deeper water, *Gabbiella humerosa*. Fifteen meters away, in a patch of emergent vegetation, *B. sudanica* (but not *B. choanomphala*) was found. *Biomphalaria choanomphala* was also occasionally recovered from the shore intermingled with *B. sudanica*, as at the Anyanga beach sampling site at Kanyibok.

**Figure 2:**
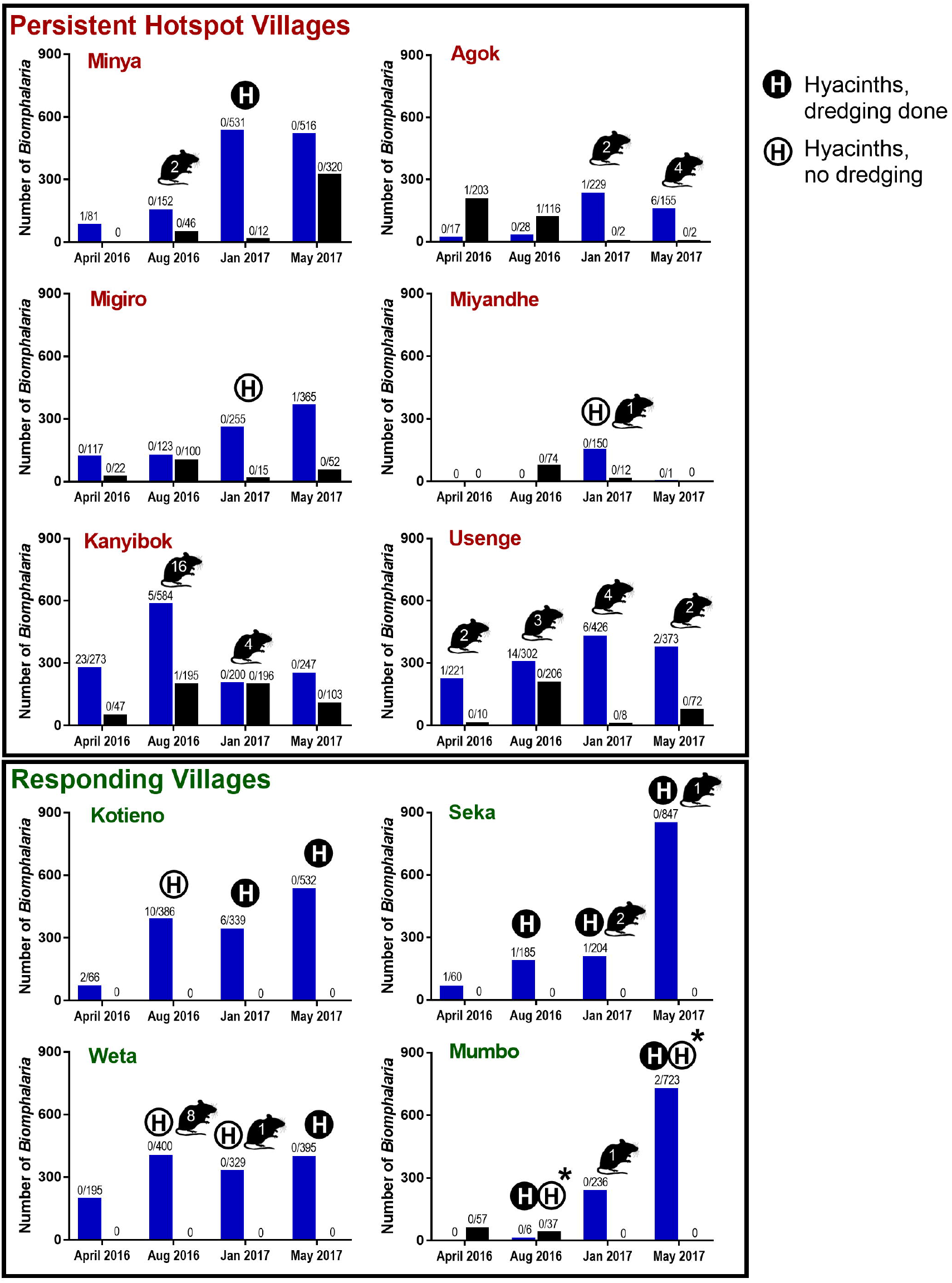
Number of *B. sudanica* (blue bars) and *B. choanomphala* (black bars) collected at each of the four sampling times (April and August 2016, January and May 2017) from the 6 persistent hotspot villages on the top versus the 4 responding villages on the bottom. Snail numbers are summed between collection sites at a village. Numbers over bars indicate number of snails infected with *S. mansoni* shedders/total number of snails found. Numbers shown within mouse figures represent the number of worms recovered from sentinel mice at that time point (summed across mice and collection sites at that village). An H in a closed circle indicates hyacinths were present and dredging was done. An H in an open circle indicates hyacinths were present but dredging could not be done. An asterisk (*) indicates Kabuong Beach was dredged but not Mumbo Beach.

**Table 2:**
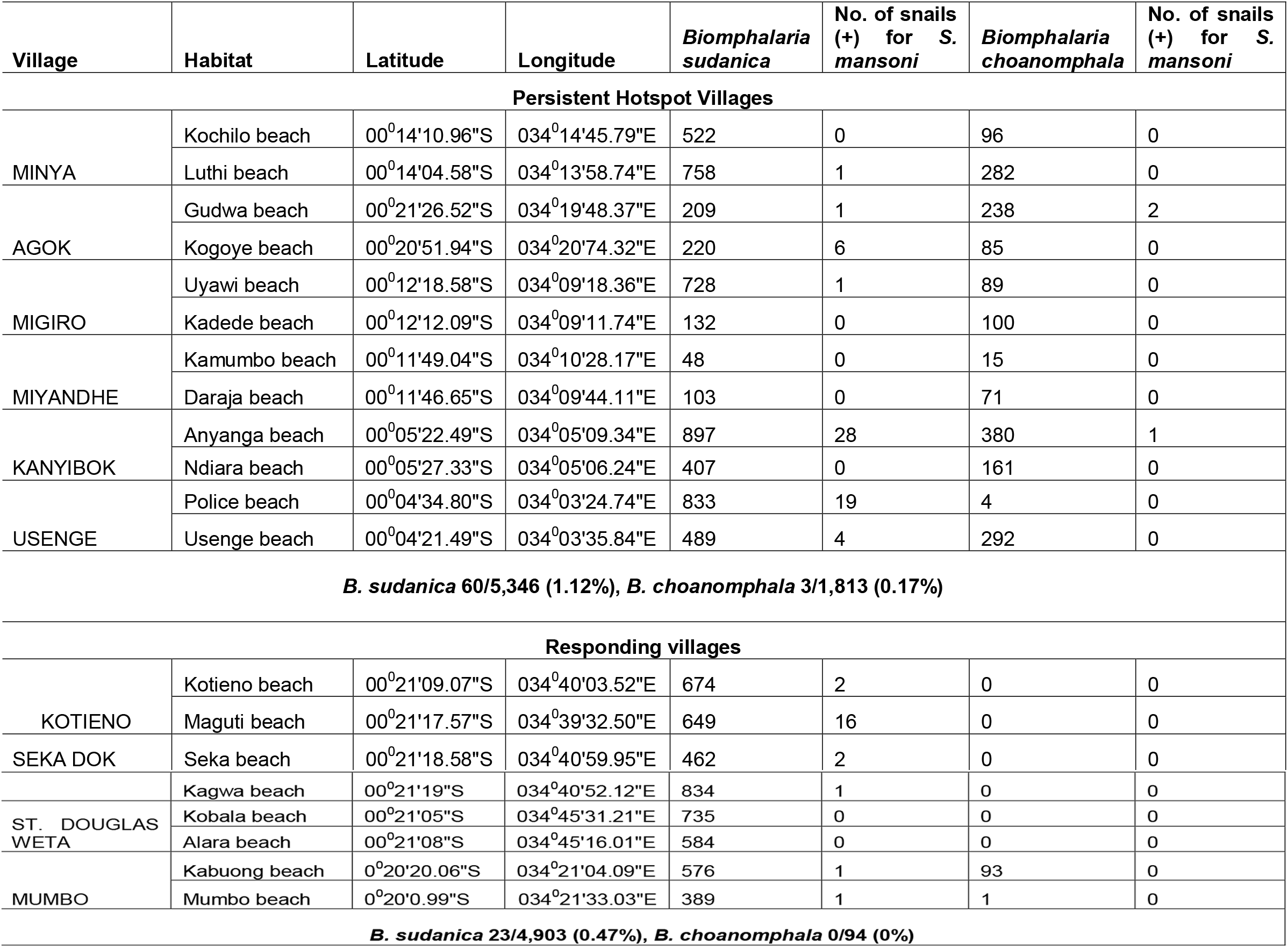
The 20 sampling sites and their GPS coordinates, total number of *Biomphalaria sudanica* and *B. choanomphala* collected from each, and the number of each species positive for *S. mansoni*.

Hyacinth mats were noted during 3 of the collective total of 48 total visits (6.25%) we paid to PHS sampling sites, and two of these times precluded dredging at such sites (Figure 1). Hyacinth mats were found at 11 of 32 collective visits (34.3%) at RESP sites, and at five of these times precluded dredging.

### Snail relative abundance

Overall, we collected far more *B. sudanica* than *B. choanomphala* (Table 2, Figure 2). However, it should be noted that because our collecting methods differed for each snail taxon, these numbers are not directly comparable (e.g. there is not necessarily a larger population of *B. sudanica* than *B. choanomphala* in the lake). The relative abundance of snails was highly variable among sites and collection times. For instance, at Miyandhe, one of the PHS villages, with its rocky, wave-swept lakeshore habitat, few snails were recovered relative to all the other villages. It was not unusual for snails to increase or decrease several fold between collection time points at the same habitats (Figure 2).

In Lake Victoria, *B. sudanica* is considered a more important intermediate host of *S. mansoni than is B. choanomphala* because *B. sudanica* with its shallow water habitat use is more likely to contact miracidia from human feces deposited along the shore. Thus, we predicted that snail driven differences between PHS and RESP sites would be reflected in this species. However, we found that PHS sites did not contain more *B. sudanica* snails than RESP sites (β = −0.0664, SE = 0.4603, P = 0.8854, Figure 3A). We note however that the three PHS villages (Minya, Kanyibok and Usenge) with the highest residual prevalence of *S. mansoni* are also the three villages with the highest overall counts of *Biomphalaria* snails. *B. sudanica* density was positively associated with the presence of hyacinths (β = 0.7704, SE = 0.2980, P = 0.0097, Figure 4) and collecting time point (β = 0.37912, SE = 0.09532, P < 0.0001).

**Figure 3:**
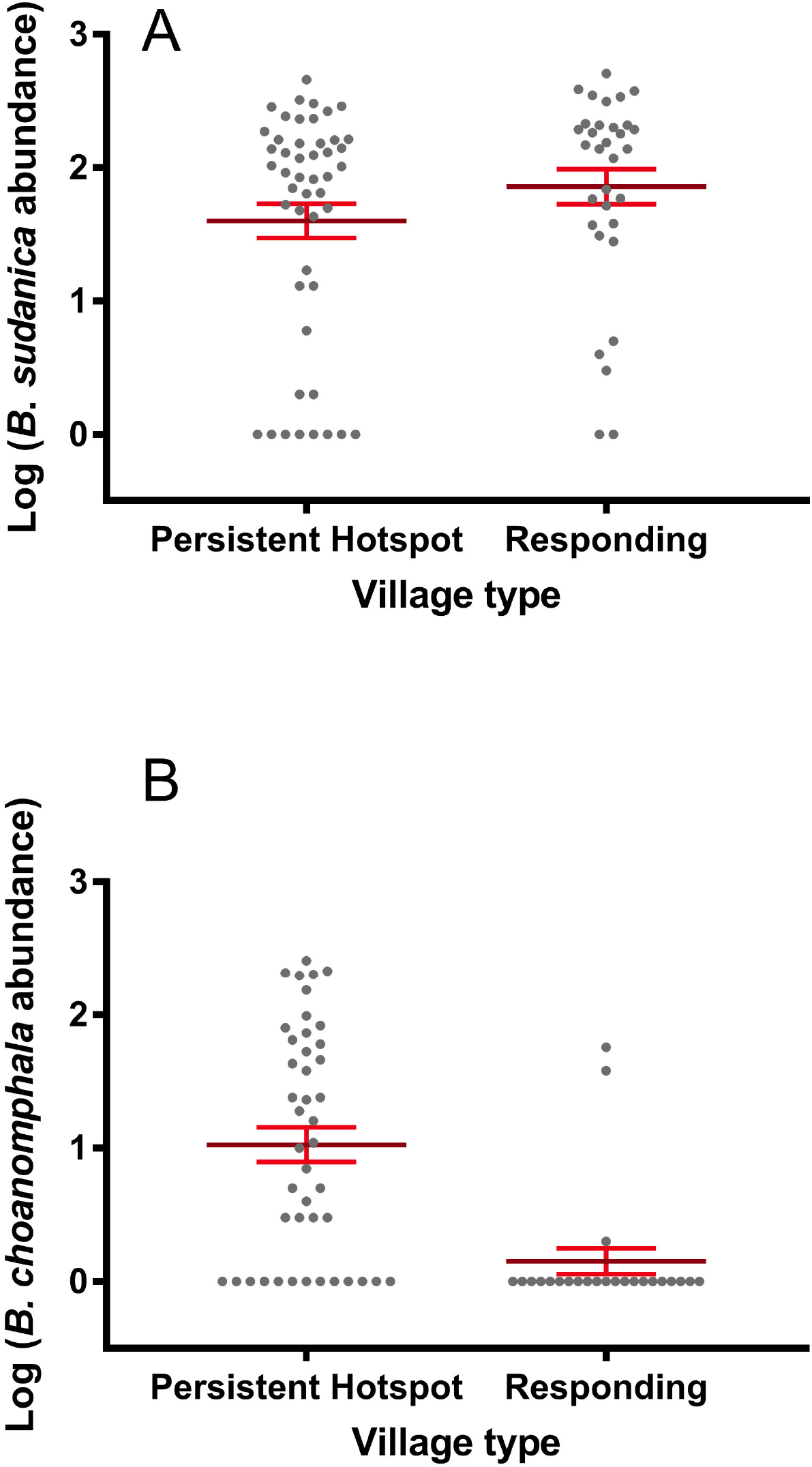
Relative abundance of *Biomphalaria* snails at persistent hotspot and responding sites of Lake Victoria in Kenya. **A.** Persistent hotspot sites did not contain more *B. sudanica* snails than responding sites when time point, village, and hyacinths were taken into account. **B.** *B. choanomphala*, was far more abundant at persistent hotspot sites compared to responding sites (P = 0.0091). Error bars indicate standard error of the mean.

**Figure 4:**
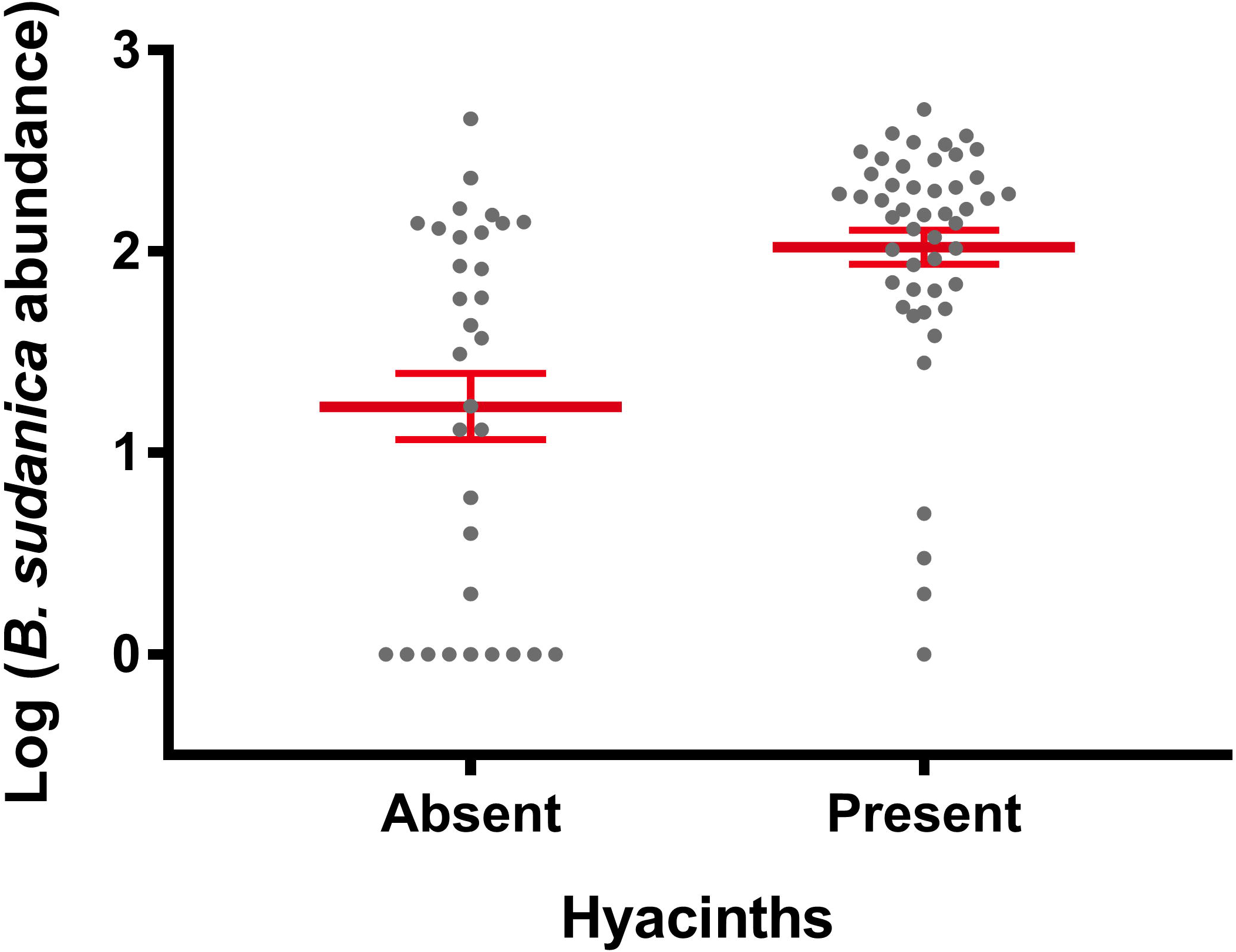
The number of *B. sudanica* collected was greater when hyacinths were present than when hyacinths were not present (P=0.00973). The presence of hyacinths did not influence *B. choanomphala* abundance (not shown). Error bars indicate standard error of the mean.

In contrast to the *B. sudanica* results, *B. choanomphala* was more abundant at PHS sites (Figure 3B) compared to RESP sites (β = −2.1644, SE = 0.6975, P = 0.0091) and the presence of hyacinths and time point were not significantly associated with *B. choanomphala* density. Thus, results of the snail analysis indicated that only the deepwater taxon, *B. choanomphala*, differed between PHS and RESP sites. The other novel finding was that *B. sudanica* abundance was positively associated with the hyacinths.

### Prevalence of infected snails

The overall number of positive snails and prevalence for *S. mansoni* recovered among all 10 villages examined were higher for *B. sudanica* (83/10,249, 0.81%) than for *B. choanomphala* (3/1,907, 0.16%). Although few *B. choanomphala* were found positive for *S. mansoni*, all came from PHS villages. For *B. sudanica*, the total number of *S. mansoni*-infected snails and overall prevalence was higher in PHS villages (60/5,346, 1.12%) than from RESP villages (23/4,903, 0.47%). As a check on the identity of schistosome cercariae recovered, we amplified the *nad5* gene from cercariae from 20 different positive snails, five from each of the time points, and representing several beach sites where schistosome cercariae were found.^28^ All 20 samples were verified as *S. mansoni* based on *nad5* band size, with one sample (Maguti Beach) exhibiting a double infection with both *S. mansoni* and *S. rodhaini*.

The distribution of *S. mansoni* infections among snails from the different villages is difficult to succinctly characterize. Two of the PHS villages with the highest residual post-treatment prevalences of *S. mansoni* in children, Kanyibok and Usenge, had the highest overall prevalence of positive snails (1.57% and 1.42% respectively (Table 2, Figure 2). From each locality though, most of these infections were found in a single collection raising the possibility they might have originated from one or a few human contamination events. Also, we found few shedders in Minya and Migoro, and none in Miyandhe, all PHS villages, whereas Kotieno, a RESP village, yielded the third most shedding snails, many of which were also found at a single time point. Ten of 24 sampling times (41.7%) from PHS villages yielded positive snails whereas 7 of 16 sampling times (43.8%) from RESP villages yielded snails positive for *S. mansoni*. Consequently, the number of *B. sudanica* infected with *S. mansoni* did not differ between PHS and RESP sites when time point, hyacinths, and village were all taken into account. There was a negative relationship that approached statistical significance between the number of infected snails collected and the presence of hyacinths at a site (β = −0.12362, SE = 0.07186, P = 0.0854), a point considered further in the discussion.

### Sentinel mice

Worm recoveries from sentinel mice (Figure 2) were low (53 worms recovered, or 0.066 worms per mouse, or 0.022 worms/mouse/hour of exposure). The greatest number of worms acquired by a single mouse was 7. Of the total number of worms recovered, 40 (75.5%) were from PHS villages (0.083 worms/mouse) while 13 (24.5%) were from RESP villages (0.041 worms/mouse). Although the total number of adult schistosomes collected from mice caged in PHS sites was greater than those caged in RESP sites, there was no statistical difference between them when village, hyacinths, and time point were considered (Figure 5, β = −0.36479 SE = 0.5720, P = 0.5237). Examination of the viscera of positive sentinel mice revealed only the presence of *S. mansoni* eggs; no *S. rodhaini* eggs were found.

**Figure 5:**
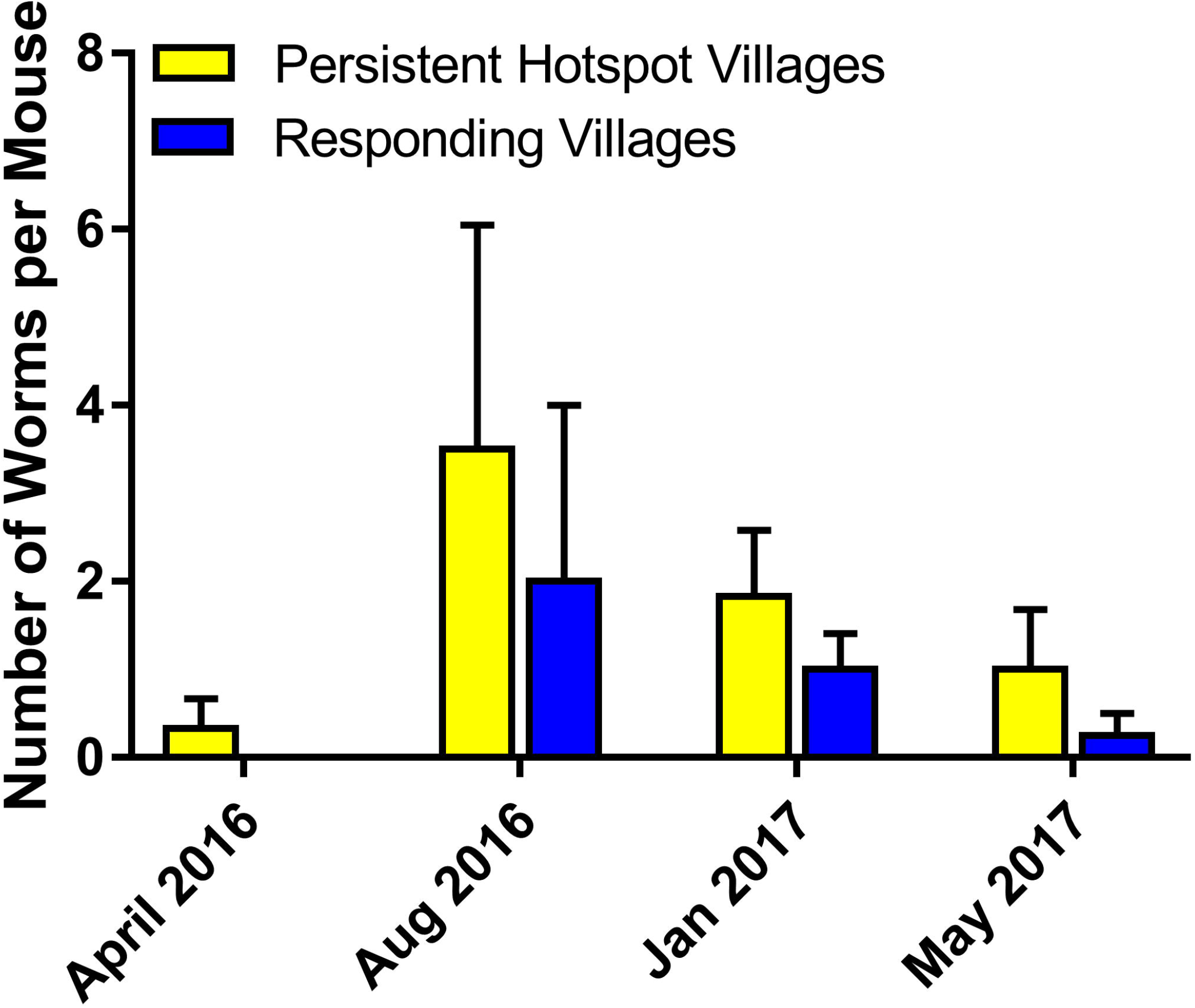
The total number of adult schistosomes collected from sentinel mice was greater in persistent hotspot than responding sites, but there was no statistical difference between them when village, hyacinths, and time point were considered.

### Correlation between snail abundance and human prevalence

Total number of *B. sudanica* collected at a village was not significantly correlated with *S. mansoni* prevalence in humans (Figure 6) either before or after the drug treatment intervention (before: Pearson r = −0. 0369, p = 0.460; after: Pearson r = −0.009, p = 0.49). However there was a strong correlation between the number of *B. choanomphala* collected at a village with *S. mansoni* prevalence in humans at the village (Figure 7) both before and after mass drug administration (before: Pearson r = 0.587, r^2^ = 0.345, p = 0.037; after: Pearson r = 0.9034, r^2^=0.816, p = 0.0002).

**Figure 6:**
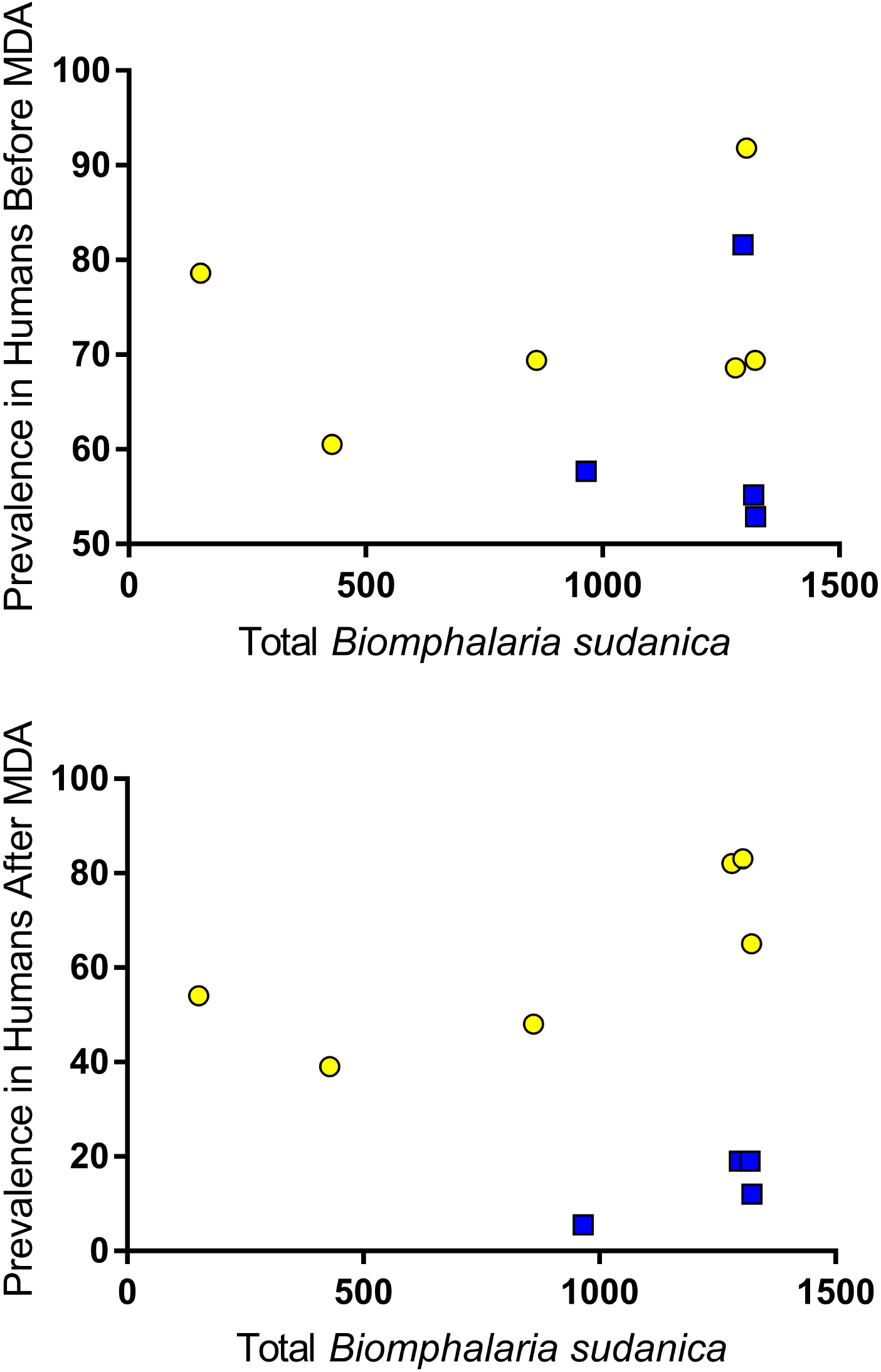
Total number of *B. sudanica* collected at a village was not significantly correlated with prevalence of *S. mansoni* in humans either before or after the drug treatment intervention (before: Pearson r = −0.0369, p = 0.460; after: Pearson r = −0.009, p = 0.49). Yellow = persistent hotspot villages; Blue = responding villages.

**Figure 7:**
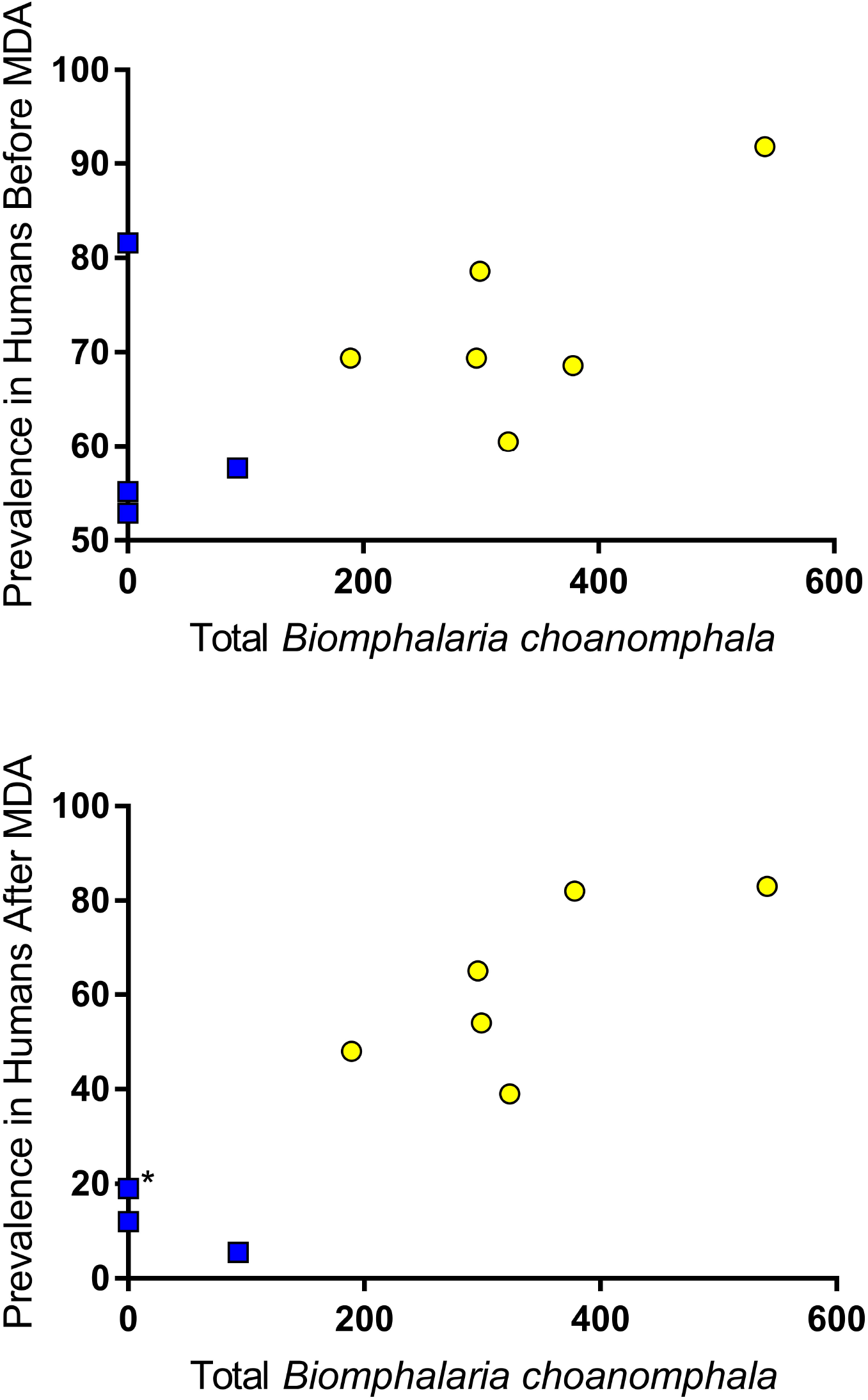
Total number of *B. choanomphala* collected at a village was significantly correlated with prevalence of *S. mansoni* in humans both before and after mass drug administration (before: Pearson r = 0.587, r^2^ = 0.345, p = 0.037; after: Pearson r = 0.9034, r^2^=0.816, p = 0.0002). Yellow = persistent hotspot villages; Blue = responding villages (* two villages with same prevalence).

### Infection of *B. sudanica* with other digenetic trematodes

In all 10 villages, the number of trematode infections other than *S. mansoni* observed from *B. sudanica* (Supplementary Table 1) exceeded those of *S. mansoni* (totals of 241 and 83, respectively). In PHS villages 60 of 184 trematode infections were of *S. mansoni* (32%), whereas in RESP villages, 23 of 150 (15%) were of *S. mansoni*. Overall prevalence of trematode infections including *S. mansoni* was similar between PHS and RESP villages (3.5 and 3.1%, respectively). Echinostomes were the most abundant non-*S. mansoni* infections with 112 snails infected from the ten study villages. We did not observe any snails to have double infections (e.g. to be simultaneously shedding two different kinds of cercariae). Although *S. mansoni* infections were consistently outnumbered by the aggregate totals of all other digenetic trematode infections, there were no statistical associations between the number of snails infected with *S. mansoni* and any of the other trematode species measured with the model employed.

With respect to having high overall total trematode prevalence, three villages stood out from the rest, the two PHS villages Kanyibok and Usenge and the one RESP village Kotieno (Supplementary Table 1). All three villages had relatively high *Biomphalaria* populations, and yielded the most *S. mansoni* infections found in snails relative to other villages (Table 2).

## DISCUSSION

The SCORE-supported PZQ-based treatment program in the Lake Victoria region in western Kenya noted that PHS villages comprised approximately a third of the 150 study village. The village prevalence was determined by testing 9-12 year old students each year. These students had a relative risk of infection that was nearly 4 times higher than for students outside the PHS area.^17^ The SCORE program is by no means unique in identifying difficulties in reducing prevalence of infection in some villages adjacent to Lake Victoria.^19^ The PHS for *S. mansoni* transmission identified by Wiegand et al., was in Siaya County, Bondo region, of western Kenya, associated with west-facing villages adjacent to the open, unprotected waters of the lake. In contrast, the PZQ RESP villages faced the more protected waters of Winam Gulf. ^17^ Among several factors postulated to be associated with the PHS we considered in our investigation were larger snail populations, or presence of different species of vector snails factors.^17^

Of the three taxa of *Biomphalaria* commonly implicated in transmission of *S. mansoni* in Kenya, two were collected in this study, the shoreline-associated taxon *B. sudanica* and the deeper water taxon *B. choanomphala*. There was no obvious indication that *B. pfeifferi* played a role in *S. mansoni* transmission in any of our study villages, as no snails of this species were found in the lake, or in streams or small impoundments within the study localities. We found *B. sudanica* at all 20 collecting sites and it was typically abundant as it is along much of the Lake Victoria shoreline.^9,11,22,34^ We recovered approximately five times as many *B. sudanica* as *B. choanomphala*.

When taking into account all sampling sites, sampling times and presence of hyacinths, mean abundance of *B. sudanica* did not differ statistically between PHS and RESP villages. We did note a significant trend for more *B. sudanica* to be present from our collecting sites if mats of hyacinths were present at the time we took our samples. Hyacinth mats were found in collecting sites from both PHS and RESP villages but were more common in the RESP villages located along Winam Gulf, an area known to be prone to hyacinth intrusions.^38^ The recovery of *B. sudanica* from hyacinths and the association we noted for them with hyacinths supports the assertion that floating hyacinth mats favor the dispersal of *B. sudanica* along the lakeshore.^35^ Hyacinth movements are likely to have major effects on the population structure of *B. sudanica* in the lake.

By contrast, *B. choanomphala* was significantly more likely to be recovered from PHS than RESP villages. We found *B. choanomphala* from all 12 of the collecting sites in PHS villages and from only 2 of the 8 collecting sites from RESP villages, both from the village of Mumbo. Mumbo is located closer to the mouth of Winam Gulf than the remaining RESP villages which are located in more protected waters of the Gulf. Previous studies have also shown *B. choanomphala* to be rare or absent within Winam Gulf, possibly because the bottom substrate is too muddy, whereas, the substrate found off the shore of the PHS villages was a combination of sand and mud, known to be favorable to *B. choanomphala*.^11,22^ Gouvras et al., found *B. choanomphala* to be focally distributed in the Tanzanian waters of the lake, including in some more protected inlets.^34^ They noted, as did we, a general trend for this species to be more common in western than eastern locations, and considered a possible gradual change in substratum conditions from west to east might be responsible. Many other factors such as levels of dissolved oxygen, pollution and seasonal effects like rainfall may also influence *B. choanomphala* populations.^33,34^ We did not observe any association between the abundance of *B. choanomphala* and water hyacinths. Although hyacinth mats can cover areas of the lake where deeper water and hence *B. choanomphala* populations are present and could have adverse shading or other effects, our impression was that hyacinth mats along the more exposed shores away from the Gulf were more prone to be dispersed by winds.

With respect to the schistosome infections among the snails recovered, a couple of caveats should be kept in mind. One is that some infections are prepatent at the time the snails are collected such that these infections would be missed. Our approach here was simply to retain all non-shedding snails in lab aquaria and to re-shed them after four weeks. This is because we have excellent snail-rearing facilities near our field sites at our disposal in which mortality of snails during the four-week interval can be minimized. This approach has the advantage of detecting those infections that successfully transition to patency (cercariae are actually produced). An alternative approach to detect prepatent infections is to employ molecular xenomonitoring techniques to detect nascent infections at the time of collection.^24,28,34,47^ Such techniques can be very sensitive and specific, but are prone to error if the infection is very young at the time of assay, and some detected infections may have been doomed to fail, so transmission potential could potentially be overestimated.^28^ Also, extractions of individual snails for PCR-based or other types of molecular detection of prepatent infections becomes prohibitively expensive and time-consuming when large numbers of snails need to be screened. Pooling of samples or use of eDNA-based techniques may help but are likely to be accompanied by a loss of precision in estimates of snail infection rates.^48^ The second caveat is that the rodent-transmitted *Schistosoma rodhaini*, the sister species of *S. mansoni*, is known to be present in the Lake Victoria basin though it is rare by comparison with *S. mansoni*.^42^ We extracted mammalian schistosome cercariae (four cercariae per infected snail) from 20 different positive snails and all yielded *nad5* bands consistent with *S. mansoni*, with one of these samples indicating a co-infection with *S. rodhaini*.^28^ Molecularly-based techniques unquestionably increase the accuracy of species identifications, but again pose some challenges insofar as an infected snail can produce thousands of cercariae over a period of several months and it is conceivable that the species composition of the schistosome cercariae could change during the interval. Again, with a large survey with many positive snails recovered, resources limit the ability to repeatedly sample individual cercariae produced by the same snail. Acknowledging this constraint, given the ubiquity of human infections in our study area, and that 20 out of 20 positive snails cercariae samples we tested were identified as *S. mansoni*, we are confident the conclusions we reach here pertain to *S. mansoni* infections.

With these caveats in mind, for snail infections of *S. mansoni*, both the number of positive snails found and prevalence were higher for *B. sudanica* than *B. choanomphala*. The overall prevalence of *S. mansoni* infections in all the *Biomphalaria* snails collected was almost 2-fold higher for the PHS villages than for the RESP villages, and the highest recovery of positive snails was found in Kanyibok (1.57%) and Usenge (1.42%), two of the villages with the highest residual prevalence of infections in schoolchildren. However, one RESP village Kotieno also showed a high overall prevalence of infected snails (1.36%). When all sampling sites, sampling times and presence of hyacinths were taken into account, PHS and RESP villages did not differ with respect to numbers or prevalence of infected *B. sudanica*. The few *B. choanomphala* found to be positive were recovered only from PHS villages.

We also noted a suggestive negative relationship between hyacinth presence and infection rates of *B. sudanica* with *S. mansoni*. If a true association, this finding may reflect a decreased likelihood of collecting infected snails when large hyacinth mats are present. Several explanations could account for this including higher mortality of infected snails due to oxygen depletion under hyacinth mats, reluctance of people to enter water for either defecation or washing purposes if hyacinths are present, or increased difficulty of miracidia in finding snails in spatially complex hyacinth mats. In general, the role of hyacinths in understanding the dynamics of schistosomiasis transmission in the lake requires further investigation.

Although fewer *B. choanomphala* than *B. sudanica* were collected, and their prevalence of infection with *S. mansoni* was relatively low, it would be incorrect to assume the former species has limited significance with respect to schistosome epidemiology in Lake Victoria. Our dredge hauls across the extensive bottom surface of the lake can only be expected to retrieve a small proportion of the snails present and likely grossly underestimate the role of this snail species in perpetuating transmission. At least two factors might facilitate infections in *B. choanomphala:* currents moving from offshore to deeper water, and fishermen who may defecate directly into deeper water from their boats. In this regard, failure to include fishermen in control programs may favor *B. choanomphala-mediated* transmission. Another factor potentially favoring involvement of *B. choanomphala* in transmission is that experimental infection studies of Kenya-derived specimens suggest this species, like *B. pfeifferi*, is inherently more susceptible to *S. mansoni* than *B. sudanica*.^29,31^ Gouvras et al., working in southern Lake Victoria found more *B. choanomphala* positive for *S. mansoni* than we did: 12.2% of all *S. mansoni* shedders were *B. choanomphala* in their study vs. only 3.5% in our study.^34^ In both studies though, *B. sudanica* was identified as the major vector for *S. mansoni* in the lake.

Considering the second measure of the force of transmission we examined, namely sentinel mice infections, we did not observe a significant difference in mean number of worms recovered between PHS and RESP villages, though worm recoveries were approximately twice as high in the former and a positive correlation was noted across villages between the total number of snails shedding *S. mansoni* and the number of worms recovered from mice. Results of snail collections and worm recoveries were concordant in 60% of our 40 sampling times, but presence of S. mansoni-shedding snails by no means guaranteed the recovery of worms from sentinel mice placed nearby, and vice versa: recovery of worms from sentinel mice was frequently noted at sampling locations and times we found no shedding snails. This suggests that neither method perfectly reflects the possibility of transmission at a particular locality at a certain time.

Not unexpectedly, and as reported from previous studies, *Biomphalaria* snails shedding other trematode cercariae were also frequently collected.^49–52^ Though non-schistosome infections exceeded *S. mansoni* infections by a factor of 2.9, we noted no significant positive or negative associations with any of the groups of trematodes with *S. mansoni* infections. The three villages that yielded the most *S. mansoni* snail infections (PHS villages Kanyibok and Usenge and RESP Kotieno) also yielded by far the most infections with other trematodes, suggesting they are general “trematode-transmitting hotspots.” One possibility to explain this is that the overall susceptibility to trematode infection is higher in snails from collecting sites in some villages than others. This possibility is currently being examined with respect to susceptibility to *S. mansoni*. Another explanation is that certain water contact points along the lakeshore attract intense activity by a variety of potential definitive hosts including people, domestic animals like cattle or goats and wild birds, the latter often thronging to locations where fishermen bring their catches. People often contact water extensively at such sites, making them potentially dangerous potential transmission sites.

In comparison to *B. sudanica, B. choanomphala* had fewer *S. mansoni* infections and fewer infections with trematodes of other species (only 4 other species noted). Deep water has been considered a refugium, or in the parlance of King et al., a “coevolutionary cold spot” that limits exposure of snails to digenetic trematodes.^52^ In comparison, shoreline habitats are visited by numerous definitive hosts and can be considered as “coevolutionary hot spots” for digenetic trematode infections, supported by the many trematode species noted to be transmitted by *B. sudanica*.^52^

This study poses several questions requiring clarification. For instance, it might be argued that PHS villages have higher prevalence of human infections because the snails adjacent to such villages are more susceptible to *S. mansoni* infection. Experiments are underway to test the relative compatibility of *B. sudanica* from different villages to sympatric and allopatric isolates of *S. mansoni*. Our null hypothesis is that strongly differentiated patterns of schistosome-snail compatibility will not be seen, because both human migration (and consequently *S. mansoni* migration) and snail movements mediated by hyacinths, winds and currents will overcome local differentiation. Also in need of additional clarification is the extent to which the gene pools of *B. sudanica* and *B. choanomphala* co-mingle. Studies of mitochondrial and nuclear markers indicate the two taxa are not highly divergent genetically and should probably be considered as a single species with the species name with precedence yet to be decided.^12,13^ It is clear though that pockets of snails with the characteristic *B. choanomphala* phenotype can be recovered from the shoreline over a period of several months. The extent to which snails of the two phenotypes interbreed, the appearance of the progeny, and the nature of genes and alleles dictating compatibility to schistosomes and other trematodes all remain to be investigated.

## CONCLUSIONS

Results of this study did not find significant differences between PHS and RESP villages for *B. sudanica* with respect to relative abundance or numbers of snails shedding *S. mansoni* cercariae, or in the number of worms recovered from sentinel mice. Although *B. sudanica* was present in all the villages we studied and based on the number of positive snails of this species recovered is also the major vector for *S. mansoni* in all villages, *B. choanomphala* was found in all PHS villages and in only one RESP village. The relative abundance of *B. choanomphala* was significantly higher in PHS villages. In our view, *B. choanomphala* may provide an under-appreciated and more diffuse and difficult to measure alternative route of transmission along much of the shore of the lake. Many other trematode species infect *Biomphalaria* snails in Lake Victoria but their overall prevalence is low and suggests they do not have a dramatic primary effect on *S. mansoni* prevalence rates in snails. Evidence of active *S. mansoni* transmission was found in all 10 villages investigated, not surprising from the standpoint that the lake supports massive populations of *Biomphalaria* snails and sanitary facilities in rural areas around the lake are scarce. However, distinct differential responsiveness to annual MDA with PZQ was somewhat surprising. Nevertheless, opportunities for transmission and reinfections are very likely to continue to occur, even in RESP villages. Control of *S. mansoni* in and around the lake will remain a daunting challenge and its persistence argues more persuasively than ever for implementation of improved sanitation and provision of safe water supplies as important parts of an integrated and sustained three-country basin-wide control approach, and as fundamental rights for the people living in the area.

## Acknowledgements

The authors thank Dr. Stephen Munga, Director, Center for Global Health Research, Kenya Medical Research Institute (KEMRI) for providing laboratory space to carry out the study, and Joseph Kinuthia, Geoffrey Maina, Polycarp Oraro and Boaz Oduor for their technical assistance in the laboratory and field sampling. Dr. Daniel G. Colley, Dr. Nupur Kittur and other members of the SCORE secretariat provided several invaluable comments. This research was undertaken with the approvals of the National Commission for Science, Technology and Innovations (permit number P/16/9609/12754) and the National Environmental Management Authority (permit number NEMA/AGR/46/2014). Field sampling of snails in the habitats was done under authorization from the Kenya Wildlife Service (permit number KWS/BRM/5001). This paper is published with the approval of the Director, KEMRI.

## Authors’ addresses

**Martin W. Mutuku**, Centre for Biotechnology Research and Development, Kenya Medical Research Institute, Nairobi, Kenya and School of Biological Sciences, College of Biological and Physical Sciences, University of Nairobi, Nairobi, Kenya. **Martina R. Laidemitt**, Center for Evolutionary and Theoretical Immunology, Parasitology Division, Museum of Southwestern Biology, Department of Biology, University of New Mexico, Albuquerque, New Mexico, United States of America. **Brianna R. Beechler**, College of Veterinary Medicine, Department of Biomedical Sciences, Oregon State University, Corvallis, Oregon, United States of America. **Ibrahim N.Mwangi**, Centre for Biotechnology Research and Development, Kenya Medical Research Institute, Nairobi, Kenya. **Fredrick. O. Otiato**, Influenza Surveillance Program, Centers for Disease Control and Prevention, Nairobi, Kenya. **Eric L. Agola**, Centre for Biotechnology Research and Development, Kenya Medical Research Institute, Nairobi, Kenya. **Horace Ochanda**, School of Biological Sciences, College of Biological and Physical Sciences, University of Nairobi, Nairobi, Kenya. **Bishoy Kamel**, Center for Evolutionary and Theoretical Immunology, Parasitology Division, Museum of Southwestern Biology, Department of Biology, University of New Mexico, Albuquerque, New Mexico, United States of America. **Gerald M. Mkoji**, Centre for Biotechnology Research and Development, Kenya Medical Research Institute, Nairobi, Kenya. **Michelle. L. Steinauer**, Department of Basic Medical Sciences, Western University of Health Sciences, Lebanon, Oregon, United States of America. **Eric S. Loker**, Center for Evolutionary and Theoretical Immunology, Parasitology Division, Museum of Southwestern Biology, Department of Biology, University of New Mexico, Albuquerque, New Mexico, United States of America.

