## Supplementary table 1 for "A search for snail-related answers to explain differences in response of *Schistosoma mansoni* to praziquantel treatment among responding and persistent hotspot villages along the Kenyan shore of Lake Victoria"

**Supplementary Table 1: Number of *B. sudanica* snails collected from each village and number of snails infected with *S. mansoni* or other trematode cercariae. SM = *Schistosoma mansoni*, ECH = echinostomes, AMPH = amphistomes, STRIG = strigeids, XIPH = xiphidiocercariae, SM:ECH = ratio of *S. mansoni* to echinostomes.**

| **V ILLAGE** | Total Snails | SM | ECH | AMPH | STRIG | XIPH | Total Number of  Infected Snails | Trematode Prevalence |
| --- | --- | --- | --- | --- | --- | --- | --- | --- |
| **Persistent Hotspot Villages** | | | | | | | | |
| MINYA | 1280 | 1 | 5 | 0 | 0 | 19 | 25 | 2.0 |
| AGOK | 429 | 7 | 4 | 0 | 1 | 7 | 19 | 4.4 |
| MIGIRO | 860 | 1 | 5 | 0 | 1 | 6 | 13 | 1.5 |
| MIYANDHE | 151 | 0 | 1 | 0 | 1 | 0 | 2 | 1.3 |
| KANYIBOK | 1304 | 28 | 22 | 7 | 4 | 1 | 62 | 4.8 |
| USENGE | 1322 | 23 | 10 | 6 | 12 | 14 | 65 | 4.9 |
| **TOTAL** | 5346 | 60 | 47 | 13 | 19 | 47 | 186 | 3.5 |
| **%** |  | 32 | 25 | 7 | 10 | 25 |  |  |
| **Responding Villages** | | | | | | | | |
| KOTIENO | 1323 | 18 | 29 | 7 | 17 | 2 | 73 | 5.5 |
| SEKA DOK | 1296 | 3 | 10 | 2 | 4 | 1 | 20 | 1.5 |
| WETA | 1319 | 0 | 13 | 1 | 6 | 15 | 35 | 2.7 |
| MUMBO | 965 | 2 | 13 | 1 | 2 | 4 | 22 | 2.3 |
| **TOTAL** | 4903 | 23 | 65 | 11 | 29 | 22 | 150 | 3.1 |
| **%** |  | 15 | 43 | 7 | 19 | 15 |  |  |
